# Biologic therapy is associated with selective changes in airway eosinophil subpopulations in severe asthma

**DOI:** 10.64898/2026.06.12.731433

**Authors:** Gabriella E. Wilson, Sandra Zaeh, Samir Gautam, Xiting Yan, Qing Liu, Olivia Hay, Nicole Grant, Jean Estrom, William W. Busse, Ruth R. Montgomery, Geoffrey L. Chupp

## Abstract

**Rationale:** Eosinophilic airway inflammation is common in severe asthma and strongly associated with symptoms, exacerbations, and impaired lung function. Although type 2 (T2)-targeted biologics improve outcomes and reduce eosinophils, many patients experience residual symptoms and exacerbations. Emerging evidence suggests that these biologics may differentially affect specific airway eosinophil subpopulations, representing a potential mechanism of suboptimal treatment response.

**Objective:** Determine the effect of biologic treatment on eosinophil subpopulations in adults with severe asthma using in-depth immune profiling with mass cytometry (CyTOF).

**Methods:** Fifty adults with severe asthma (28 biologic-naïve, 22 on stable-dose biologic therapy for ≥6 months) underwent clinical phenotyping, spirometry, blood sampling, and sputum induction. Twenty-nine sputum samples passed quality control thresholds and were profiled by CyTOF. Manually gated sputum eosinophils were clustered using FlowSOM to identify eosinophil subpopulations, and cluster abundances and marker expression were compared across treatment groups.

**Measurements and Main Results:** CyTOF revealed treatment-associated shifts in circulating immune cells (lower CD4+ T cells and B cells, higher monocytes) and lower sputum CD8+ T cells. Unsupervised clustering of sputum eosinophils identified eight distinct subpopulations, and selective depletion of Cluster 6 was noted in biologic-treated participants (biologic-naïve vs anti-TSLP logFC −4.98, p=0.003; biologic-naïve vs anti-IL5 logFC −6.89, p=0.01). Higher Cluster 6 proportion correlated with worse ACT scores (rho = −0.44, p = 0.02) and lung function (FEV1 % predicted: rho = −0.47, p < 0.01; FEV1/FVC: rho = −0.40, p = 0.03). Functionally, Cluster 6 displayed enriched trafficking/activation markers (CCR3/Eotaxin-1, CD69, CD80, CRTH2) and non-T2 inflammatory mediators (TNFα, IL-8, TLR7).

**Conclusion:** Biologic therapy in severe asthma was associated with selective depletion of a highly activated sputum eosinophil subpopulation with capability to drive both T2 and non-T2 inflammatory pathways. This cluster correlated with worse asthma control and lung function, indicating it may be a biologically important driver of persistent disease and potential biomarker to more accurately predict treatment response.

## Introduction

Asthma is a highly prevalent, heterogenous chronic inflammatory airway disease that affects approximately 300 million people worldwide.^1^ About 10% of patients with asthma experience severe disease that is refractory to standard therapies, exhibiting persistent symptoms, airflow limitation, and recurrent exacerbations that contribute to reduced quality of life and high healthcare utilization.^2, 3^ Most patients with severe asthma exhibit a Type 2 (T2) inflammatory signature characterized by accumulation of eosinophils in the airways, but the response of these patients to T2 targeted biologics is highly variable. This underscores the need for a comprehensive understanding of the pathways of eosinophilic inflammation for both biomarker development, targeted therapeutic strategies, and identification of novel mechanisms that contribute to severe asthma.^4, 5^

Eosinophils have long been recognized as an important effector cell in asthma with clear contributions to several hallmarks of the disease, including epithelial damage, airway hyperresponsiveness, and mucus hypersecretion.^6^ As a biomarker, blood and sputum eosinophil counts have been associated with reduced lung function, high symptom burden, and increased exacerbation risk.^7–9^ Sputum eosinophils, in particular, provide a direct measurement of airway inflammation and strongly correlate with disease activity and exacerbation risk.^10^ Over the last 15 years, several biologic therapies targeting T2 inflammation have emerged, including monoclonal antibodies directed against interleukin-5 (IL-5), IL-5 receptor α, interleukin-4 receptor α (IL-4Rα), and thymic stromal lymphopoietin (TSLP). These have transformed the management of severe asthma, with clear efficacy at reducing exacerbations, improving lung function, and facilitating weaning of maintenance oral corticosteroids.^11–14^ Many of these biologics have also been shown to reduce blood eosinophils and, to a lesser extent, sputum eosinophils.^12, 15–17^ Despite this relative eosinophil depletion, however, a subset of patients continues to experience persistent symptoms and exacerbations despite treatment with biologic therapy, highlighting our incomplete understanding of the roles of eosinophils in asthma and the specific effects that biologic medications have on eosinophils in the airway.

Apart from their roles as effector cells, eosinophils also carry out several homeostatic functions across tissues, including supporting tissue modeling/repair, metabolic regulation in adipose tissue, and maintenance of mucosal immune homeostasis.^18–22^ This functional diversity has led to increased interest in eosinophil subpopulations, and growing evidence suggests that specific eosinophil subpopulations may carry out distinct functional roles in health and disease. Additionally, distinct eosinophil subpopulations may respond differently to T2 biologics, which represents an important potential mechanism for suboptimal response to these advanced therapies. In this regard, we recently performed a sub-study of the MUPPITS-2 clinical trial which examined sputum eosinophils from children with severe asthma using mass cytometry (CyTOF).^23^ This work demonstrated that a subset of sputum eosinophils with intermediate expression of CD62L exhibited an activated phenotype and was enriched in children receiving mepolizumab who experienced breakthrough exacerbations compared with those who remained exacerbation-free during the study. These findings suggest that treatment-resistant eosinophil populations within the airway may contribute to ongoing disease manifestations and underscore the need for a deeper understanding of airway eosinophil heterogeneity and the contributions of distinct airway eosinophil subpopulations to asthma pathogenesis. Furthermore, examining residual airway eosinophils and other airway immune cells in patients treated with biologics provides a unique opportunity to uncover critical insights into mechanisms of disease, identify biomarkers of response to treatment, and reveal novel therapeutic targets.

## Methods

### Study design

The study protocol and informed consent materials were reviewed by the Yale Institutional Review Board prior to study initiation and the study was registered in the Research Registry. Adult patients with severe asthma were recruited through the Yale Center for Asthma and Airways Disease at the Winchester Center for Lung Disease for participation in the study from April 2024 – October 2025. Patients had to be either biologic-naïve (defined as no prior biologic for 6 months prior to enrollment) with plans to initiate biologic treatment (as decided by their primary pulmonologist), or biologic treated (defined as being on stable-dose biologic for ≥6 months prior to enrollment). Full inclusion and exclusion criteria for the DILIGENT study are shown in the supplemental material (Table S1). Informed consent was obtained, and consented participants underwent medical history, pulmonary function testing, blood collection by phlebotomy, and sputum induction in the initial study visit (V1).^24^ Biologic-naïve participants started biologic treatment within 1 week of their initial study visit. All participants presented for repeat sampling at a second study visit conducted 16 weeks following V1.

### Sample Processing, Eosinophil isolation by CyTOF, and Eosinophil clustering

Sample processing was performed immediately after collection, as previously described.^25^ Sputum cells were assessed microscopically, and samples that met quality standards (total cell count ≥0.5×10^6^ cells, viability ≥50%, and squamous cell percentage ≤20%) were processed for CyTOF analysis. For participants who were unable to produce a high-quality sputum sample, buffy coat was isolated from whole blood samples and processed for CyTOF. Surface staining for CyTOF was performed for cell lineage surface markers prior to fixation. ^25–27^ Samples were then stored at −80C and thawed in batches for intracellular staining and CyTOF processing. A full list of surface and intracellular antibodies used are shown in Table S2.

Bead normalization was conducted in Matlab using Nolan Lab Normalization, and normalized files were uploaded to CytoBank, where manual gating of sputum cells was performed, as previously described.^23^ Manually gated sputum eosinophils were then computationally clustered using FlowSOM implemented in the CATALYST framework in R (Bioconductor). Clustering was performed using an eosinophil-specific marker panel (Table S3) and repeated using the full marker panel to assess for stability in the eosinophil clusters identified (Table S2). Single-cell data was visualized using UMAP based on the selected type markers.^23^ To characterize cluster phenotypes, heatmaps of median marker expression were generated using the ComplexHeatmap package. For each cluster, median expression values were calculated across all cells and scaled from 0 to 1 for each marker to facilitate comparison across clusters.

### Data Analysis

Baseline clinical characteristics, age, and BMI were compared between treatment groups using Mann Whitney test, and sex and race/ethnicity were compared using Chi-sq test (Table 1). Relative abundance of manually gated immune cells in the airway and blood were analyzed as the proportion of leukocytes, and comparisons were made between Biologic-naïve and Biologic-treated participants with Mann-Whitney test.

|  | Biologic Naïve<br>(n=28) | Biologic Treated<br>(n=22) | p-value* |
| --- | --- | --- | --- |
| Demographics, *mean (SD) or count (%) unless specified* |  |  |  |
| Age | 51 (15) | 63 (15.8) | <0.01 |
| Sex |  |  |  |
| Female | 19 (68) | 12 (55) | 0.34 |
| Male | 9 (32) | 10 (45) |  |
| Race/Ethnicity |  |  |  |
| Black | 10 (36) | 3 (14) | 0.09 |
| White | 14 (50) | 18 (82) | 0.09 |
| Latino | 6 (21) | 1 (0.5) | 0.09 |
| BMI | 33.2 (11.3) | 31.8 (11.1) | 0.3 |
| Clinical Characteristics, *mean (SD) or median (IQR) unless specified* |  |  |  |
| Exacerbations† | 1.5 (0-3) | 0 (0-1) | <0.01 |
| ACT score | 16 (12-20) | 21 (17-23) | <0.01 |
| Blood Eosinophils | 250 (123-425) | 70 (40-180) | <0.01 |
| Total IgE# | 195 (45-577) | 96 (42-333) | 0.4 |
| FEV1/FVC | 0.66 (0.13) | 0.65 (0.16) | 0.87 |
| FEV1 (% pred) | 73 (21) | 81 (30) | 0.3 |
| Positive BDR§ | 7 (25) | 2(9) |  |
| FeNO (ppb) | 40 (32) | 35 (23) | 0.87 |
| Definition of Abbreviations: BMI = body mass index, FEV1 = forced expiratory volume in 1 second, FVC = forced vital capacity, pred = percent predicted, FeNO = fraction of exhaled nitric oxide, BDR = positive bronchodilator response (+12% and 200cc) |  |  |  |
| * Mann-Whitney Test performed for Age, BMI, Exacerbations, ACT Score, Blood eosinophils, IgE, FEV1/FVC, FEV1, and FENO |  |  |  |
| Chi-sq Test performed for Sex, Race/Ethnicity, and positive BDR |  |  |  |
| † Within the preceding 12 months prior to enrollment |  |  |  |
| # Values reflect most recent historical IgE |  |  |  |
| § Count (%) shown |  |  |  |

Differences in sample-level median marker expression was calculated across all manually gated eosinophils and compared between treatment groups (biologic-naïve vs biologic treated) using Wilcoxon rank-sum test.

Differential abundance (DA) analysis of eosinophil clusters between treatment groups was measured using the diffcyt framework with edgeR-based statistical testing to compare cluster proportions across samples (version 4.8.2). Library sizes were normalized using the trimmed mean of M-values (TMM) method. Differential abundance was evaluated using quasi-likelihood (QL) negative binomial generalized linear models with Bayes moderation of dispersion estimates. A three-group design matrix was constructed to compare biologic-naïve, anti-TSLP-treated, and anti-IL5-treated subjects, and pairwise comparisons were evaluated between groups. Log fold-change (logFC) values represent the log2-transformed difference in normalized cluster cell counts between groups. Given the limited number of prespecified cluster comparisons, nominal p-values were used to assess statistical significance.

Eosinophil clusters with significant abundance differences between treatment groups were focused on for further functional analyses. Associations between abundance of eosinophil clusters of interest and clinical characteristics were assessed using Spearman rank correlation. To define the phenotypic features of eosinophil clusters of interest, expression was summarized within each sample as the median arcsinh-transformed expression among the eosinophils in the cluster of interest and among all remaining eosinophils. These within-sample paired summaries were compared using the Wilcoxon signed-rank test, and p values were adjusted for multiple testing using the Benjamini-Hochberg false discovery rate method. Effect sizes are reported as the median paired difference in expression.

## Results

50 participants were enrolled in the study and underwent detailed immune profiling of sputum (n=29) or blood (n=21). Of the 50 participants enrolled, 28 participants were biologic naïve at enrollment, and 22 were on stable dose biologic for at least 6 months (tezepelumab n=11, dupilumab n=6, benralizumab n=2, mepolizumab n=2, and omalizumab, n=1). Biologic-treated participants were older than biologic-naïve individuals (mean age 61 vs 51, p<0.01), had significantly lower blood eosinophil counts (116 vs 303, p<0.01) and significantly higher asthma control test (ACT) scores (20 vs 16, p<0.01). Additional demographic and clinical characteristics are shown in Table 1.

Blood and sputum immune cell abundance was assessed by CyTOF via manual gating of all samples and compared between biologic-naïve and biologic-treated participants (Figure 1). Biologic treated participants exhibited lower circulating CD4+ T cells (28% lower [95% CI 2.55-13.65], p=0.006) and B cells (34% lower [95% CI 0.86-5.57], p=0.01), and higher monocytes (46% lower [95% CI 4.20-14.40], p=0.0003), as shown in Figure 1A. Biologic treated participants also displayed significantly reduced sputum CD8+ T cells compared to biologic-naïve participants (58% lower [95% CI 0.03-1.02], p=0.04), as shown in Figure 1B.

**Figure 1:**
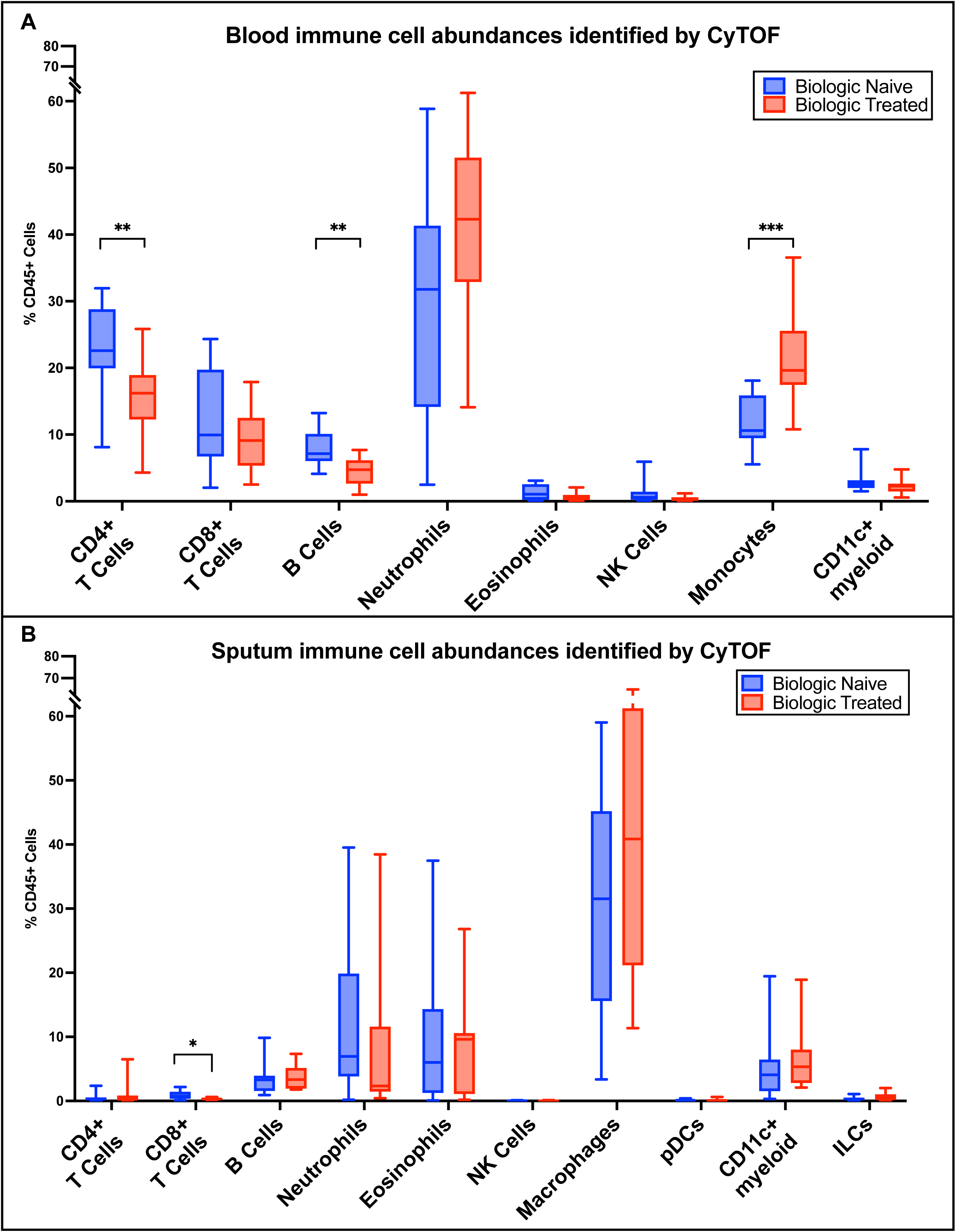
Relative abundances of sputum and blood leukocytes according to biologic treatment status. **A**, Manually gated sputum leukocytes are shown as a proportion of total CD45+ cells in biologic-naïve and biologic-treated participants. **B**, Manually gated blood leukocytes are shown as a proportion of total CD45+ cells in biologic-naïve and biologic-treated participants. Individual leukocyte proportions per sample and median and interquartile range for each cell type are shown.

**Figure 2:**
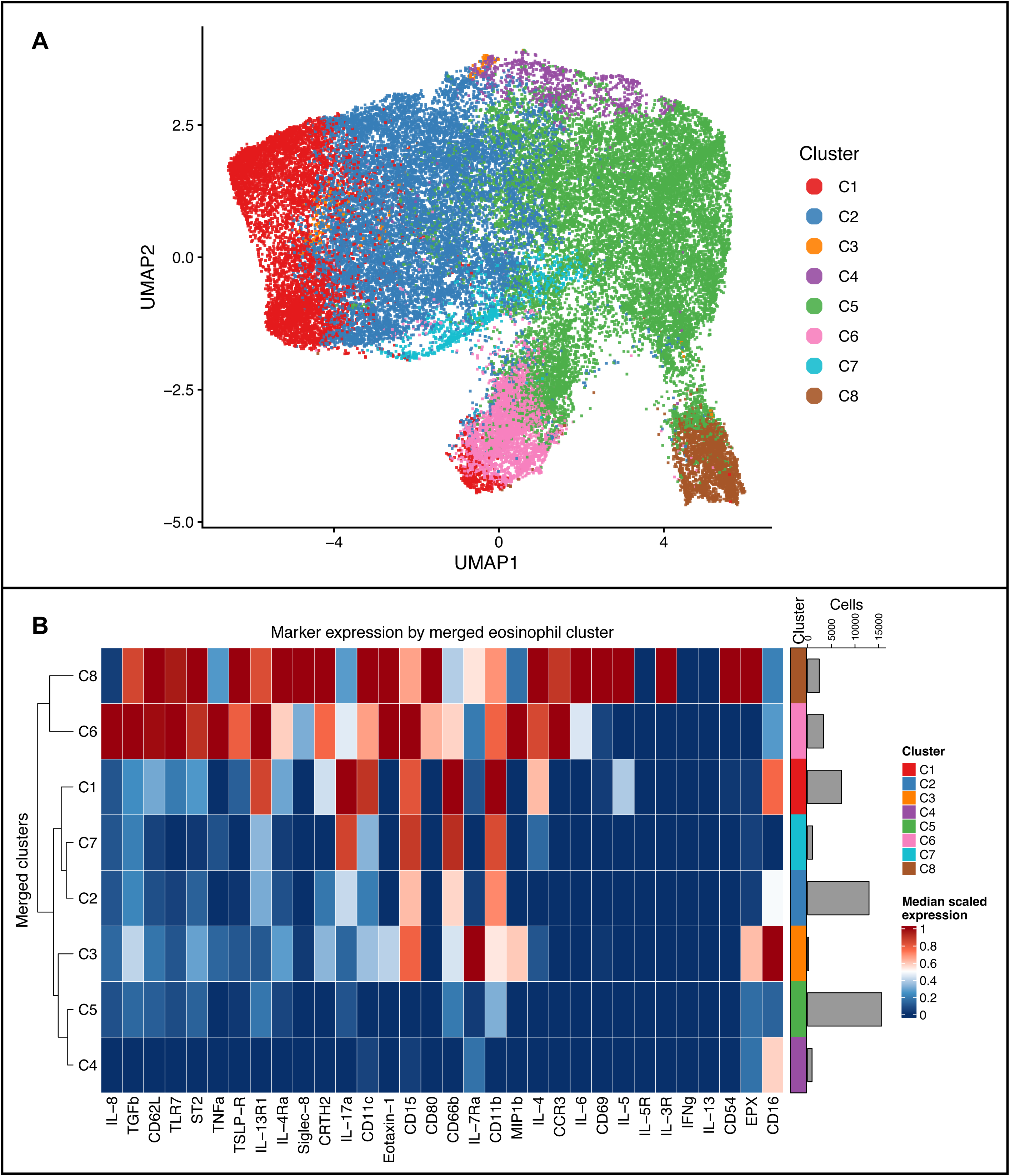
Clustering of manually gated sputum eosinophils reveals 8 distinct subpopulations of eosinophils. **A,** UMAP representation of 8 distinct sputum eosinophil subpopulations identified. **B,** Median scaled expression of eosinophil functional markers across the 8 eosinophil subpopulations identified. Inter-cluster similarity is shown on the dendrogram on the left of the heatmap. Clustering was performed on manually gated sputum eosinophils isolated from biologic-naïve and biologic treated patients, and markers used for clustering are shown in Table S3.

Unsupervised clustering of manually gated eosinophils identified 10 clusters of eosinophils. Two pairs of clusters demonstrated substantial overlap in both marker expression profiles and UMAP localization, consistent with redundant phenotypes. These clusters were therefore merged, yielding eight distinct eosinophil clusters. UMAP visualization demonstrated regional segregation of these clusters with minimal overlap, supporting the presence of phenotypically distinct eosinophil subpopulations (Figure 3A). This was further supported by heatmap analysis of median marker expression, which revealed distinct expression profiles across clusters (Figure 3B). Overall and per-sample counts and proportions for each eosinophil cluster is shown in supplemental Tables S4 and S5, respectively.

**Figure 3:**
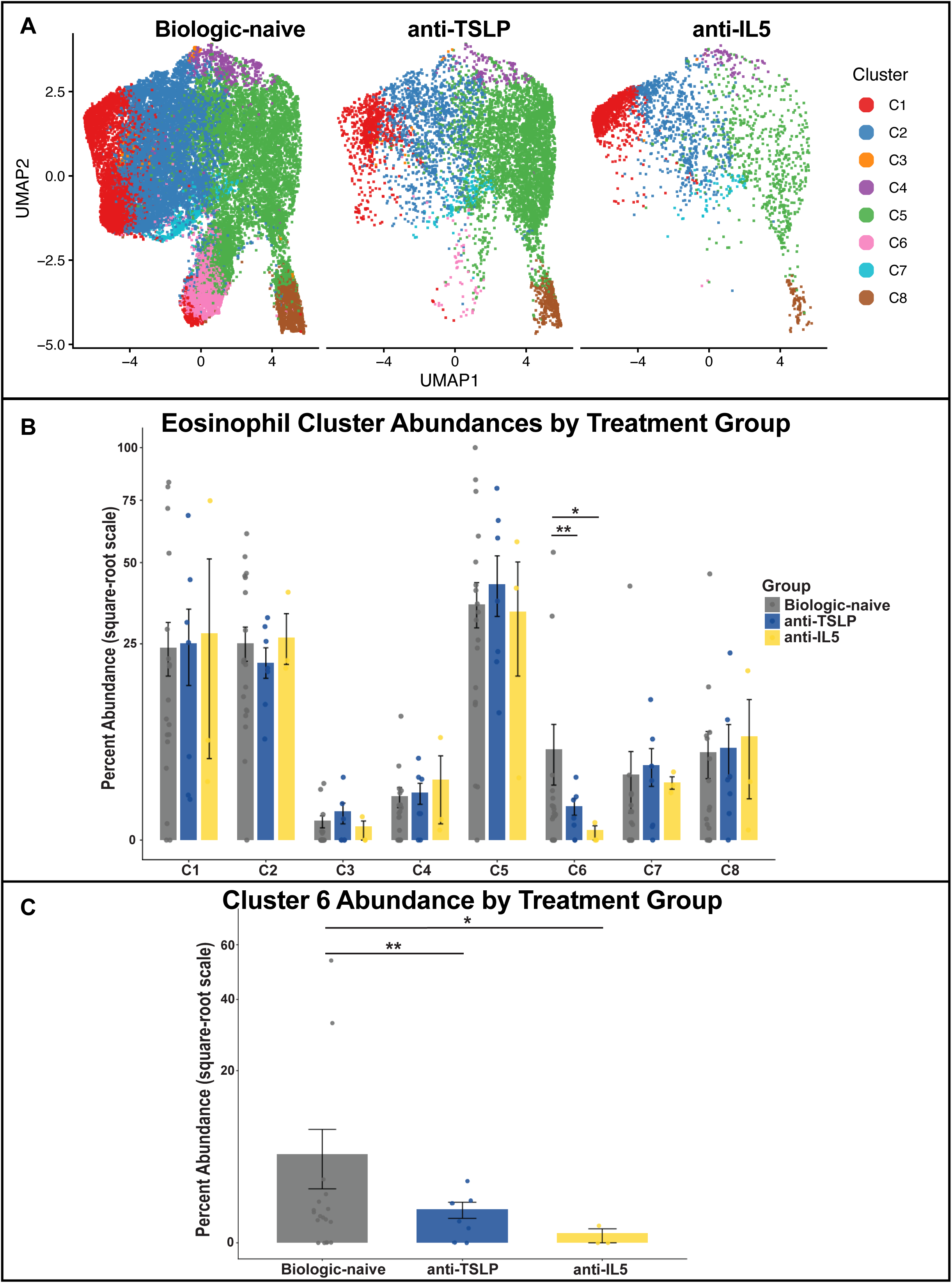
Relative abundance of eosinophil clusters across biologic-naïve, anti-TSLP, and anti-IL5 treated subjects. Data shown for all clusters by UMAP (A) and bar graph (B); data shown for cluster 6 abundance by bar graph (C). Bars represent mean percent abundance per sample, with overlaid points indicating individual subjects and error bars denoting standard error of the mean (SEM). The y-axis is displayed on a square-root scale to improve visualization of low-abundance populations.

Differential abundance analysis of the merged eosinophil clusters revealed that Cluster 6 was significantly reduced in biologic-treated participants compared with biologic-naive individuals (Figure 4). This effect was observed in both anti-TSLP–treated subjects and anti-IL5–treated participants (logFC = −4.98, p=0.003 and logFC = −6.89, p=0.01, respectively), and there was no difference in cluster 6 abundance between anti-TSLP and anti-IL5 treated participants. Besides Cluster 6, no other eosinophil clusters demonstrated significant differences in abundance across treatment groups. Furthermore, increased proportion of C6 eosinophils was associated with worse lung function and asthma control, including inverse correlations with FEV1 (% predicted, rho = −0.47, p < 0.01), FEV1/FVC (rho = −0.40, p = 0.03), and ACT score (rho = −0.44, p = 0.02) as shown in Figure S1.

**Figure 4:**
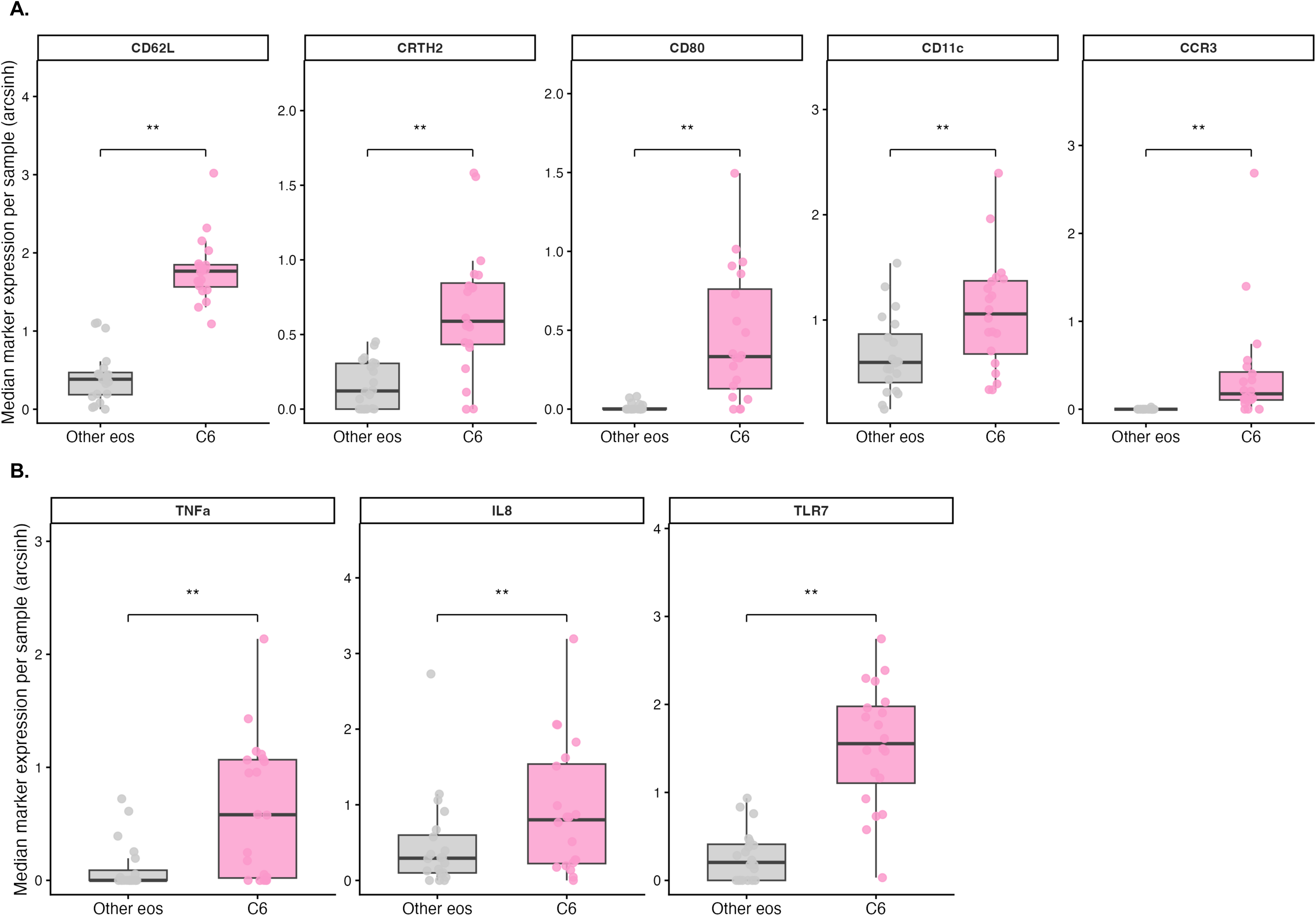
Inflammatory marker expression in Cluster 6 compared to all other sputum eosinophils. Boxplots show sample-level median expression in cluster 6 compared with all other eosinophil clusters pooled. Each point represents a single sample. Center lines indicate medians, boxes represent interquartile ranges (IQR), and whiskers extend to 1.5× IQR. Functional markers are separated into (A) eosinophil activation and trafficking markers, and (B) innate and non-T2 inflammatory mediators.

Cluster 6 eosinophils (which represented about 7.7% of all eosinophils identified) demonstrated a distinct expression profile compared to the other clusters, with significantly higher expression of several eosinophil trafficking and activation markers when compared to all other eosinophils, including higher L-selectin (CD62L, median difference 1.36, FDR=0.001), CRTH2 (median difference 0.43, FDR=0.002), CD80 (median difference 0.32, FDR=0.002), CD11c (median difference 0.29, FDR=0.006), and CCR3 (median difference 0.17, FDR=0.002), (Figure 4A). Furthermore, Cluster 6 was noted to express high levels of several inflammatory mediators and receptors typically attributed to non-T2 mechanisms of disease pathogenesis in asthma, including TNFα (median difference 0.45, FDR=0.004), IL-8 (median difference 0.47, FDR=0.003), and TLR7 (median difference 1.38, FDR=0.001) as shown in Figure 4B. Interestingly, when comparing cluster 6 to each eosinophil cluster identified, TNFα and IL-8 and were the top 2 enriched markers discriminating cluster 6 from the other eosinophil clusters identified when ranked based on the number of significant comparisons and consistency of expression differences across clusters. These findings indicate that Cluster 6 has potential to drive both T2 and non-T2 mediated inflammatory pathways. Clusters 6 also exhibited higher levels of alarmin receptor expression compared to the other eosinophils, indicating that this cluster may have increased capacity to drive airway inflammation in response to epithelial-derived signals (Figure S2). A heatmap demonstrating the top 15 markers enriched in Cluster 6 compared to the other eosinophil clusters identified is shown in Figure S3. Individual pairwise comparisons of cluster 6 with each other eosinophil cluster demonstrated similar findings (Figure S4). LogFC and FDR for pairwise comparisons of T2, non-T2, and eosinophil activation and trafficking marker expression in C6 compared with the other eosinophils clusters identified is shown in supplemental Table S6.

## Discussion

In this severe asthma cohort, biologic-treated participants demonstrated significant differences compared to biologic naïve participants across multiple domains, including baseline clinical characteristics, circulating and sputum immune cell populations, and sputum eosinophil subpopulation abundances. We found that biologic-treated participants had significantly higher ACT scores, fewer exacerbations in the prior year, and reduced blood eosinophil levels compared to biologic-naïve participants. Biologic therapy was associated with broad shifts in circulating lymphoid and myeloid cells and reduced sputum CD8+ T cells, but the dominant finding associated with biologic treatment was a selective depletion of one phenotypically distinct eosinophil subpopulation (Cluster 6) rather than equal reduction in sputum eosinophil subpopulations. This finding suggests that asthma biologics may have distinct effects on specific eosinophil subpopulations, which may be relevant to clinical response to these treatments. Furthermore, the similar findings noted between anti-TSLP and anti-IL5 treated participants in this study suggest that Cluster 6 eosinophils are sensitive to both IL-5 and alarmin mediated inflammatory pathways.

Cluster 6 eosinophils exhibit higher expression of eosinophil trafficking and activation markers (CCR3/Eotaxin-1 axis, CD80, CRTH2), suggesting recent recruitment and activation in the setting of airway inflammation, as well as potentially enhanced immunostimulatory potential^28, 29^. Cluster 6 eosinophils also exhibit high levels of CD62L, a commonly used marker for identifying eosinophil subpopulations^22, 30–32^. We previously identified 3 subpopulations of sputum eosinophils in children which exhibited low, intermediate and high levels of CD62L expression (termed CD62L^lo^, CD62L^int^, and CD62L^hi^ eosinophils). In that study, CD62L^int^ eosinophils exhibited the most activated phenotype and were higher in children with breakthrough exacerbations on mepolizumab, suggesting that this activated eosinophil subpopulation may be an important driver of treatment response.^23, 33^ Here we found substantial overlap in marker expression between Cluster 6 eosinophils and the CD62L^int^ eosinophil subpopulation identified in the MUPPITS2 cohort, suggesting that these eosinophil subpopulations are analogous. Additionally, our current work demonstrates a clear negative correlation between Cluster 6 sputum eosinophils and asthma control test (ACT) scores and physiologic impairment (FEV1 percent predicted, and FEV1/FVC), further supporting the notion that eosinophil subpopulations may have distinct roles that impact clinical manifestations of disease.

In addition to exhibiting a highly activated phenotype, the Cluster 6 eosinophil subpopulation identified here was also enriched for several markers linked to non-T2 inflammatory pathways, including IL-8 and TNFα. This expression profile suggests that this subpopulation of airway eosinophils may participate more broadly in driving airway inflammation across the T2-non-T2 axis, including engagement of neutrophil-associated pathways via IL-8 and TNFα.^34–36^ With such broad inflammatory potential, inadequate suppression or re-emergence of this subpopulation may underscore incomplete clinical response to treatment. Finally, enrichment for these non-T2 inflammatory mediators was also noted in the CD62L^int^ eosinophils identified in the MUPPITS2 cohort, further supporting the analogous nature between these two sputum eosinophil subpopulations identified in adults and children with asthma.

There are several limitations that should be considered when interpreting our findings. First, our sample size resulted in limited power for subgroup analyses by biologic class, and although 50 participants were enrolled, only 29 were able to produce an adequate sputum sample, which may reflect deeper, non-random differences in those participants compared to participants who were unable to produce sputum. Additionally, although our Cluster 6 eosinophil subpopulation was reproducibly identified across multiple analyses, independent validation and functional studies are needed to define its biological significance. Finally, the cross-sectional nature of this study limits our ability to establish a causal relationship between biologic treatment and depletion of Cluster 6 eosinophils. Future studies in larger, multicenter cohorts with longitudinal sampling before and after biologic treatment are needed to validate these findings and determine whether Cluster 6 eosinophils can serve as biomarkers of treatment response.

In conclusion, these findings suggest that CD62L-expressing sputum eosinophils reflect an activated subpopulation that may have potential as a biomarker for biologic response. This will be further assessed with ongoing work that will examine the longitudinal stability of these eosinophil subpopulations, and whether depletion of this Cluster 6 eosinophil subpopulation with biologic treatment correlates with clinical response to treatment and exacerbation risk. The combined enrichment for T2-associated features and non-T2 inflammatory mediators support the concept that airway eosinophils should be evaluated not only by abundance, but also by immunophenotype, so that we can more precisely understand their roles as biomarkers and targets in adults with asthma.

