## Supplemental Material for "Biologic therapy is associated with selective changes in airway eosinophil subpopulations in severe asthma"

**Table S1:** Inclusion/Exclusion Criteria

| Inclusion Criteria | Exclusion Criteria |
| --- | --- |
| 1. Ages 18 to 85 years 2. A clinician diagnosis of severe asthma on medium-to-high dose ICS/LABA that is a biologic candidate due to inadequate control or already taking a biologic to maintain control 3. Historical evidence of variable airflow obstruction determined by variability in FEV1 between visits or an increase in FEV1 after an inhaled beta-agonist, inhaled corticosteroids, or an oral prednisone burst 4. On stable-dose biologic (³6 months prior to enrollment) OR Biologic naïve (no biologic treatment for asthma within 6 months prior to enrollment) and being initiated on biologic treatment for severe asthma by their pulmonary provider 5. Willing to undergo study procedures | 1. >20 pack-year smoking history 2. Active smoking 3. Uncontrolled severe chronic conditions (such as congestive heart failure, renal failure, liver disease, chronic viral infections) 4. Conditions that limit ability to safely undergo the study procedures required for participation 5. Pre-existing clinically important lung condition other than asthma 6. Systemic maintenance oral corticosteroid (OCS) use or recent oral corticosteroid burst within 4 weeks of enrollment 7. Fever or evidence of an active infection 8. Investigator concerns about participant’s willingness to adhere to study requirements |

**Table S2:** CyTOF Antibody Panel

| Gating Markers | | | Surface Markers | | Intracellular Markers | |
| --- | --- | --- | --- | --- | --- | --- |
| Metal Isotope | Marker | Gate | Metal Isotope | Marker | Metal Isotope | Marker |
| 89Y | CD45 | Spike-in control | 141Pr | CD124/IL4Ra | 143Nd | IL5 |
| 110Cd | CD45 | Leukocytes | 142Nd | CD11b | 148Nd | Eotaxin/Ccl11 |
| 111Cd | CD4 | CD4+ T cell | 144Nd | CD69 | 150Nd | MIP1b |
| 112Cd | CD8a | CD8+ T cell | 145Nd | CD125/IL5R | 152Sm | TNFa |
| 114Cd | CD20 | B cells | 147Sm | ST2 | 154Sm | TGF-Beta1 |
| 146Nd | CD294/CRTH2 | ILC2 | 151Eu | CD123/IL3R | 156Gd | IL6 |
| 149Sm | CD127/IL7Ra | ILC | 155Gd | TSLPR | 163Dy | IL4 |
| 153Eu | BDCA2/CD303 | pDC | 161Dy | CD80 | 168Er | IFNg |
| 158Gd | EPX | Eosinophils | 166Er | IL13R1 | 169Tm | IL13 |
| 159Tb | CD11c | mDC | 167Er | CD287/TLR7 | 172Yb | IL17A |
| 160Gd | CD14 | Macrophage | 170Er | CD54 | 173Yb | IL8 |
| 162Dy | cytokeratin | Epithelial cells | 174Yb | CD62L |  |  |
| 164Dy | CD15 | Granulocytes | 175Lu | CD193/CCR3 |  |  |
| 165Ho | Siglec-8 | Eosinophils |  |  |  |  |
| 171Yb | CD66b | Granulocytes |  |  |  |  |
| 176Yb | CD56 | NK Cells |  |  |  |  |
| 209Bi | CD16 | Neutrophils |  |  |  |  |

**Table S3:** Eosinophil Clustering Marker Panel

| Metal | Marker |
| --- | --- |
| 142-Nd | CD11b |
| 144-Nd | CD69 |
| 145-Nd | IL-5R |
| 151-Eu | IL-3R |
| 152-Sm | TNFa |
| 156-Gd | IL-6 |
| 158-Gd | EPX |
| 161-Dy | CD80 |
| 163-Dy | IL-4 |
| 164-Dy | CD15 |
| 165-Ho | Siglec-8 |
| 168-Er | IFNg |
| 169-Tm | IL-13 |
| 170-Er | CD54 |
| 171-Yb | CD66b |
| 174-Yb | CD62L |
| 175-Lu | CCR3 |
| 209-Bi | CD16 |

**Table S4:** Eosinophil cluster abundances

| **Cluster** | **Number of cells** | **Percent proportion** |
| --- | --- | --- |
| **C1** | 7151 | 16.3% |
| **C2** | 12966 | 29.6% |
| **C3** | 230 | 0.5% |
| **C4** | 919 | 2.1% |
| **C5** | 15663 | 35.8% |
| **C6** | 3363 | 7.7% |
| **C7** | 1043 | 2.4% |
| **C8** | 2450 | 5.6% |

**Table S5:** Per-sample eosinophil cluster cell counts and proportions

| **Sample ID** | **Biologic Status** | **Cluster** | **Cluster Cell Count** | **Total Cell Count** | **Percent of Total** |
| --- | --- | --- | --- | --- | --- |
| DIL002.1S | Biologic Naïve | C1 | 78 | 146 | 53.42 |
|  |  | C2 | 30 | 146 | 20.55 |
|  |  | C4 | 1 | 146 | 0.68 |
|  |  | C5 | 18 | 146 | 12.33 |
|  |  | C6 | 4 | 146 | 2.74 |
|  |  | C7 | 4 | 146 | 2.74 |
|  |  | C8 | 11 | 146 | 7.53 |
| DIL005.1S | Biologic Naïve | C1 | 498 | 599 | 83.14 |
|  |  | C2 | 80 | 599 | 13.36 |
|  |  | C4 | 2 | 599 | 0.33 |
|  |  | C5 | 11 | 599 | 1.84 |
|  |  | C7 | 8 | 599 | 1.34 |
| DIL009.1S | Biologic Naïve | C1 | 116 | 1612 | 7.20 |
|  |  | C2 | 177 | 1612 | 10.98 |
|  |  | C4 | 13 | 1612 | 0.81 |
|  |  | C5 | 507 | 1612 | 31.45 |
|  |  | C6 | 10 | 1612 | 0.62 |
|  |  | C7 | 675 | 1612 | 41.87 |
|  |  | C8 | 114 | 1612 | 7.07 |
| DIL011.1S | Biologic Naïve | C1 | 384 | 4033 | 9.52 |
|  |  | C2 | 1604 | 4033 | 39.77 |
|  |  | C3 | 2 | 4033 | 0.05 |
|  |  | C4 | 32 | 4033 | 0.79 |
|  |  | C5 | 1702 | 4033 | 42.20 |
|  |  | C6 | 31 | 4033 | 0.77 |
|  |  | C7 | 8 | 4033 | 0.20 |
|  |  | C8 | 270 | 4033 | 6.69 |
| DIL013.1S | Biologic Naïve | C2 | 4 | 19 | 21.05 |
|  |  | C5 | 15 | 19 | 78.95 |
| DIL017.1S | Biologic Naïve | C5 | 43 | 43 | 100.00 |
| DIL019.1S | Biologic Naïve | C1 | 1 | 567 | 0.18 |
|  |  | C2 | 57 | 567 | 10.05 |
|  |  | C4 | 9 | 567 | 1.59 |
|  |  | C5 | 478 | 567 | 84.30 |
|  |  | C6 | 9 | 567 | 1.59 |
|  |  | C8 | 13 | 567 | 2.29 |
| DIL020.1S | Biologic Naïve | C1 | 158 | 2189 | 7.22 |
|  |  | C2 | 1334 | 2189 | 60.94 |
|  |  | C3 | 1 | 2189 | 0.05 |
|  |  | C4 | 78 | 2189 | 3.56 |
|  |  | C5 | 569 | 2189 | 25.99 |
|  |  | C6 | 16 | 2189 | 0.73 |
|  |  | C7 | 31 | 2189 | 1.42 |
|  |  | C8 | 2 | 2189 | 0.09 |
| DIL022.1S | Biologic Naïve | C1 | 50 | 234 | 21.37 |
|  |  | C2 | 122 | 234 | 52.14 |
|  |  | C3 | 1 | 234 | 0.43 |
|  |  | C4 | 4 | 234 | 1.71 |
|  |  | C5 | 56 | 234 | 23.93 |
|  |  | C6 | 1 | 234 | 0.43 |
| DIL030.1S | Biologic Naïve | C1 | 1 | 5 | 20.00 |
|  |  | C2 | 1 | 5 | 20.00 |
|  |  | C5 | 3 | 5 | 60.00 |
| DIL031.1S | Biologic Naïve | C1 | 129 | 3849 | 3.35 |
|  |  | C2 | 320 | 3849 | 8.31 |
|  |  | C5 | 1299 | 3849 | 33.75 |
|  |  | C6 | 2070 | 3849 | 53.78 |
|  |  | C7 | 20 | 3849 | 0.52 |
|  |  | C8 | 11 | 3849 | 0.29 |
| DIL033.1S | Biologic Naïve | C1 | 278 | 3212 | 8.66 |
|  |  | C2 | 129 | 3212 | 4.02 |
|  |  | C3 | 1 | 3212 | 0.03 |
|  |  | C4 | 2 | 3212 | 0.06 |
|  |  | C5 | 1613 | 3212 | 50.22 |
|  |  | C6 | 1046 | 3212 | 32.57 |
|  |  | C7 | 5 | 3212 | 0.16 |
|  |  | C8 | 138 | 3212 | 4.30 |
| DIL035.1S | Biologic Naïve | C1 | 1516 | 7735 | 19.60 |
|  |  | C2 | 3505 | 7735 | 45.31 |
|  |  | C3 | 129 | 7735 | 1.67 |
|  |  | C4 | 111 | 7735 | 1.44 |
|  |  | C5 | 1167 | 7735 | 15.09 |
|  |  | C6 | 88 | 7735 | 1.14 |
|  |  | C7 | 41 | 7735 | 0.53 |
|  |  | C8 | 1178 | 7735 | 15.23 |
| DIL038.1S | Biologic Naïve | C1 | 238 | 1868 | 12.74 |
|  |  | C2 | 836 | 1868 | 44.75 |
|  |  | C4 | 10 | 1868 | 0.54 |
|  |  | C5 | 756 | 1868 | 40.47 |
|  |  | C6 | 7 | 1868 | 0.37 |
|  |  | C7 | 10 | 1868 | 0.54 |
|  |  | C8 | 11 | 1868 | 0.59 |
| DIL040.1S | Biologic Naïve | C1 | 5 | 7 | 71.43 |
|  |  | C2 | 2 | 7 | 28.57 |
| DIL047.1S | Biologic Naïve | C1 | 18 | 287 | 6.27 |
|  |  | C2 | 31 | 287 | 10.80 |
|  |  | C4 | 1 | 287 | 0.35 |
|  |  | C5 | 104 | 287 | 36.24 |
|  |  | C6 | 1 | 287 | 0.35 |
|  |  | C8 | 132 | 287 | 45.99 |
| DIL059.1S | Biologic Naïve | C1 | 901 | 3121 | 28.87 |
|  |  | C2 | 1442 | 3121 | 46.20 |
|  |  | C3 | 65 | 3121 | 2.08 |
|  |  | C4 | 311 | 3121 | 9.96 |
|  |  | C5 | 371 | 3121 | 11.89 |
|  |  | C6 | 9 | 3121 | 0.29 |
|  |  | C8 | 22 | 3121 | 0.70 |
| DIL060.1S | Biologic Naïve | C1 | 512 | 632 | 81.01 |
|  |  | C2 | 96 | 632 | 15.19 |
|  |  | C4 | 3 | 632 | 0.47 |
|  |  | C5 | 12 | 632 | 1.90 |
|  |  | C6 | 3 | 632 | 0.47 |
|  |  | C7 | 5 | 632 | 0.79 |
|  |  | C8 | 1 | 632 | 0.16 |
| DIL004.1S | Anti-TSLP | C1 | 18 | 1372 | 1.31 |
|  |  | C2 | 91 | 1372 | 6.63 |
|  |  | C3 | 4 | 1372 | 0.29 |
|  |  | C4 | 33 | 1372 | 2.41 |
|  |  | C5 | 910 | 1372 | 66.33 |
|  |  | C6 | 2 | 1372 | 0.15 |
|  |  | C7 | 2 | 1372 | 0.15 |
|  |  | C8 | 312 | 1372 | 22.74 |
| DIL016.1S | Anti-TSLP | C1 | 647 | 2552 | 25.35 |
|  |  | C2 | 755 | 2552 | 29.58 |
|  |  | C3 | 22 | 2552 | 0.86 |
|  |  | C4 | 111 | 2552 | 4.35 |
|  |  | C5 | 946 | 2552 | 37.07 |
|  |  | C6 | 8 | 2552 | 0.31 |
|  |  | C7 | 3 | 2552 | 0.12 |
|  |  | C8 | 60 | 2552 | 2.35 |
| DIL034.1S | Anti-TSLP | C1 | 12 | 39 | 30.77 |
|  |  | C2 | 10 | 39 | 25.64 |
|  |  | C3 | 1 | 39 | 2.56 |
|  |  | C4 | 1 | 39 | 2.56 |
|  |  | C5 | 9 | 39 | 23.08 |
|  |  | C6 | 1 | 39 | 2.56 |
|  |  | C7 | 5 | 39 | 12.82 |
| DIL041.1S | Anti-TSLP | C1 | 26 | 38 | 68.42 |
|  |  | C2 | 7 | 38 | 18.42 |
|  |  | C5 | 4 | 38 | 10.53 |
|  |  | C8 | 1 | 38 | 2.63 |
| DIL048.1S | Anti-TSLP | C1 | 13 | 287 | 4.53 |
|  |  | C2 | 55 | 287 | 19.16 |
|  |  | C5 | 170 | 287 | 59.23 |
|  |  | C6 | 3 | 287 | 1.05 |
|  |  | C7 | 19 | 287 | 6.62 |
|  |  | C8 | 27 | 287 | 9.41 |
| DIL053.1S | Anti-TSLP | C1 | 38 | 3515 | 1.08 |
|  |  | C2 | 420 | 3515 | 11.95 |
|  |  | C4 | 16 | 3515 | 0.46 |
|  |  | C5 | 2824 | 3515 | 80.34 |
|  |  | C6 | 43 | 3515 | 1.22 |
|  |  | C7 | 125 | 3515 | 3.56 |
|  |  | C8 | 49 | 3515 | 1.39 |
| DIL057.1S | Anti-TSLP | C1 | 96 | 218 | 44.04 |
|  |  | C2 | 70 | 218 | 32.11 |
|  |  | C4 | 1 | 218 | 0.46 |
|  |  | C5 | 45 | 218 | 20.64 |
|  |  | C7 | 5 | 218 | 2.29 |
|  |  | C8 | 1 | 218 | 0.46 |
| DIL007.1S | Anti-IL5R | C1 | 1129 | 1510 | 74.77 |
|  |  | C2 | 316 | 1510 | 20.93 |
|  |  | C4 | 1 | 1510 | 0.07 |
|  |  | C5 | 38 | 1510 | 2.52 |
|  |  | C6 | 3 | 1510 | 0.20 |
|  |  | C7 | 22 | 1510 | 1.46 |
|  |  | C8 | 1 | 1510 | 0.07 |
| DIL036.1S | Anti-IL5R | C1 | 8 | 360 | 2.22 |
|  |  | C2 | 69 | 360 | 19.17 |
|  |  | C4 | 1 | 360 | 0.28 |
|  |  | C5 | 208 | 360 | 57.78 |
|  |  | C7 | 7 | 360 | 1.94 |
|  |  | C8 | 67 | 360 | 18.61 |
| DIL058.1S | Anti-IL5 | C1 | 73 | 1128 | 6.47 |
|  |  | C2 | 450 | 1128 | 39.89 |
|  |  | C3 | 4 | 1128 | 0.35 |
|  |  | C4 | 77 | 1128 | 6.83 |
|  |  | C5 | 465 | 1128 | 41.22 |
|  |  | C7 | 34 | 1128 | 3.01 |
|  |  | C8 | 25 | 1128 | 2.22 |
| DIL001.1S | Anti-IgE | C1 | 208 | 2608 | 7.98 |
|  |  | C2 | 953 | 2608 | 36.54 |
|  |  | C4 | 101 | 2608 | 3.87 |
|  |  | C5 | 1320 | 2608 | 50.61 |
|  |  | C6 | 8 | 2608 | 0.31 |
|  |  | C7 | 14 | 2608 | 0.54 |
|  |  | C8 | 4 | 2608 | 0.15 |

**Table S6:** LogFC and FDR values for pairwise comparisons of functional marker expression between Cluster 6 (C6) and the other eosinophil clusters identified

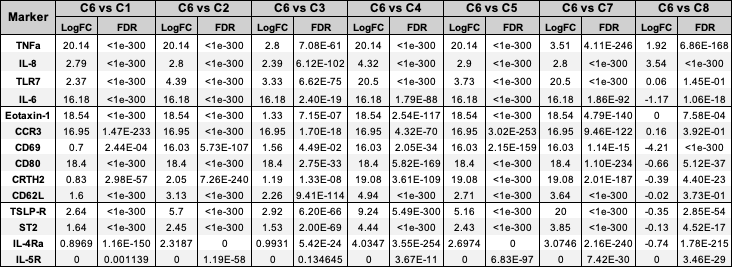

**Supplemental Figure Legends:**

**Figure S1: Correlation between Cluster 6 proportion and clinical characteristics of disease.** Spearman correlation shown for correlation testing between Cluster 6 and (A) Asthma control test (ACT) score, (B) FEV1 (% predicted), and (C) FEV1/FVC.

**Figure S2: Alarmin receptor marker expression across eosinophil subtypes.** Boxplots show sample-level median expression in cluster 6 compared with all other eosinophil clusters pooled. Each point represents a single sample. Center lines indicate medians, boxes represent interquartile ranges (IQR) and whiskers extend to 1.5× IQR.

**Figure S3: Functional marker expression comparison between Cluster 6 and all remaining eosinophil clusters (B).** Heatmap showing log2 fold-change in marker expression for cluster 6 relative to each of the remaining eosinophil clusters (C1–C5, C7–C8). Values represent log2 fold-change calculated from median expression levels, with positive values (red) indicating higher expression in cluster 6 and negative values (blue) indicating lower expression. Each cell displays the log2 fold-change and corresponding false discovery rate (FDR) derived from Wilcoxon rank-sum testing with Benjamini–Hochberg correction.

**Figure S4: Functional marker expression between Cluster 6 and each other eosinophil cluster identified.**

Boxplots show sample-level median expression within each cluster. Center lines indicate medians, boxes represent interquartile ranges (IQR) and whiskers extend to 1.5× IQR. Each point represents the median expression value for a given sample within the indicated cluster. Functional markers are separated into (A) eosinophil activation and trafficking markers, and (B) innate and non-T2 inflammatory mediators, and (C) Alarmin receptor expression.

**Figure S1**:

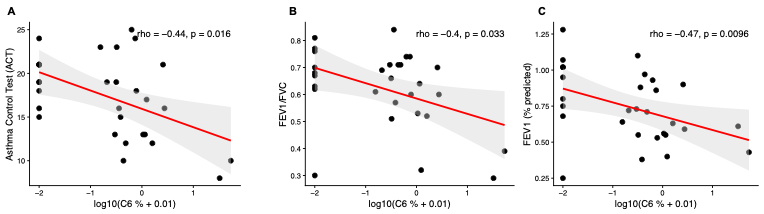

**Figure S2:**

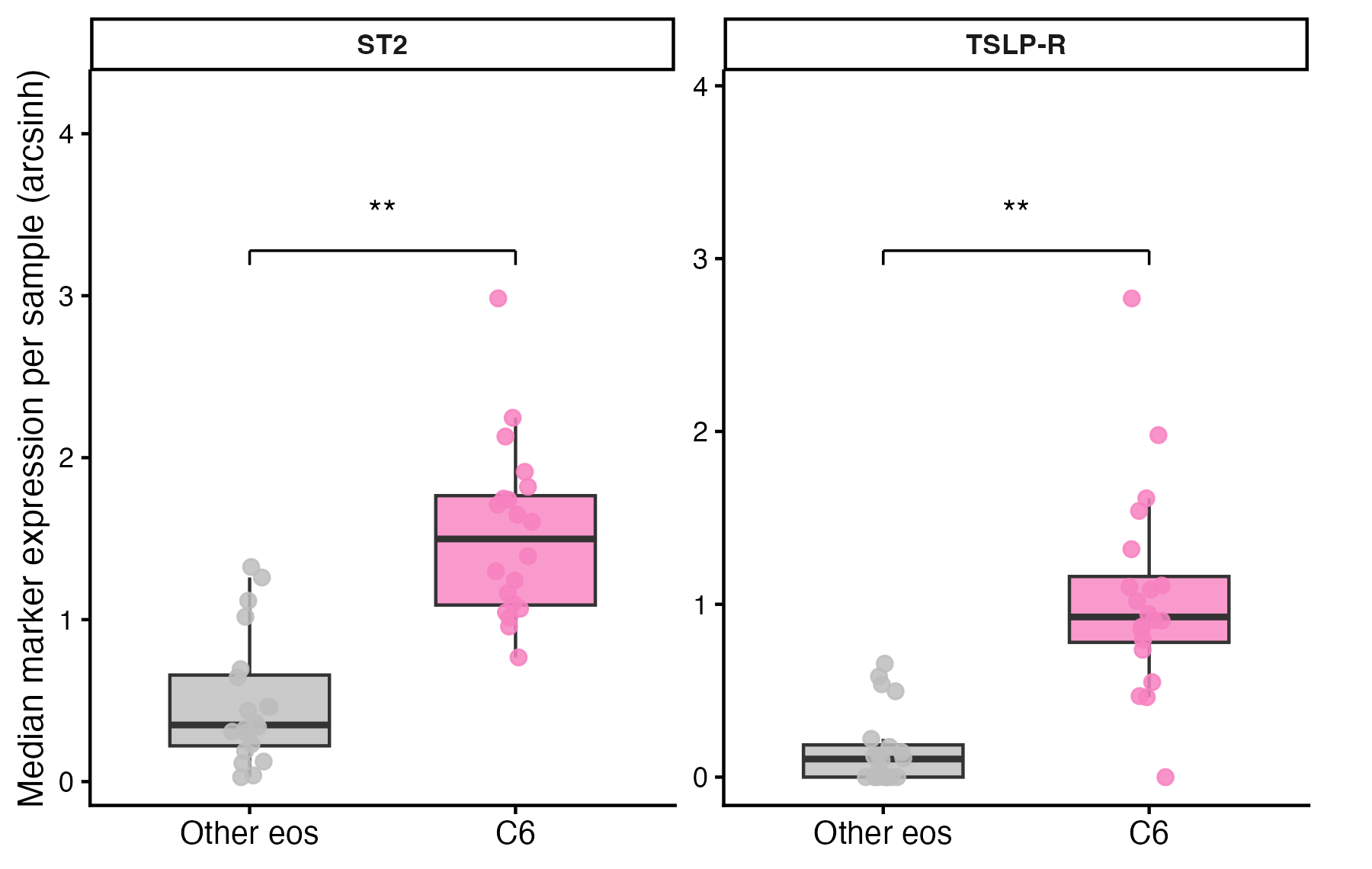

**Figure S3:**

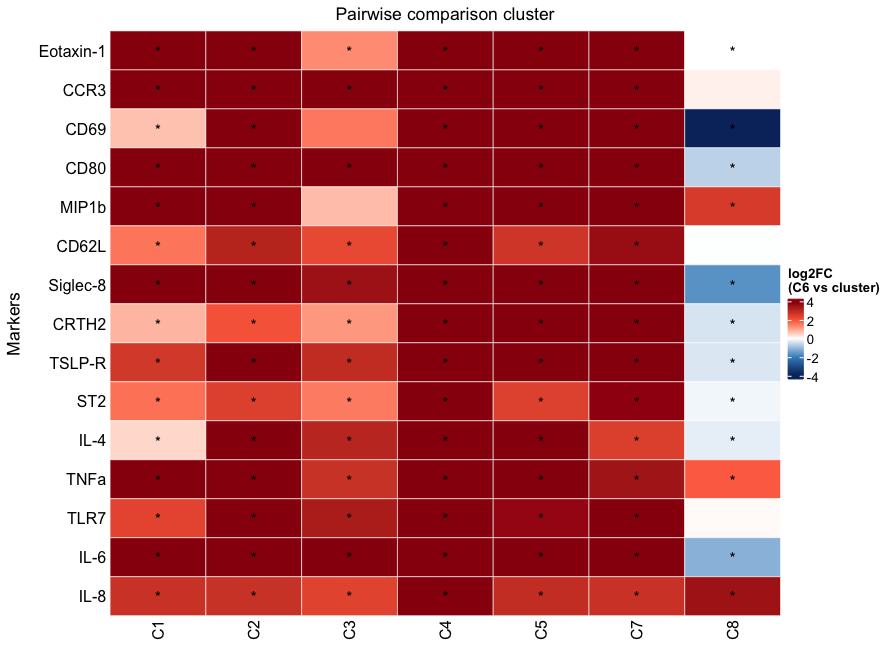

**Figure S4:**

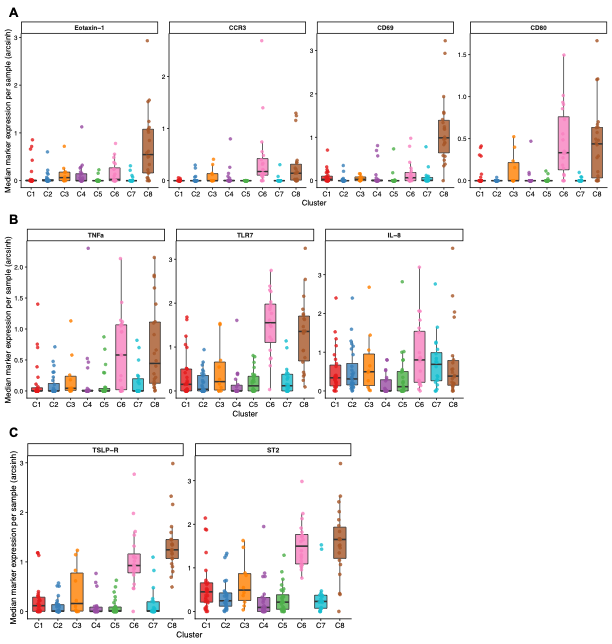
